## supplemental figures and tables for "AdRoit: an accurate and robust method to infer complex transcriptome composition"

### Affiliations

### Supplementary Figures

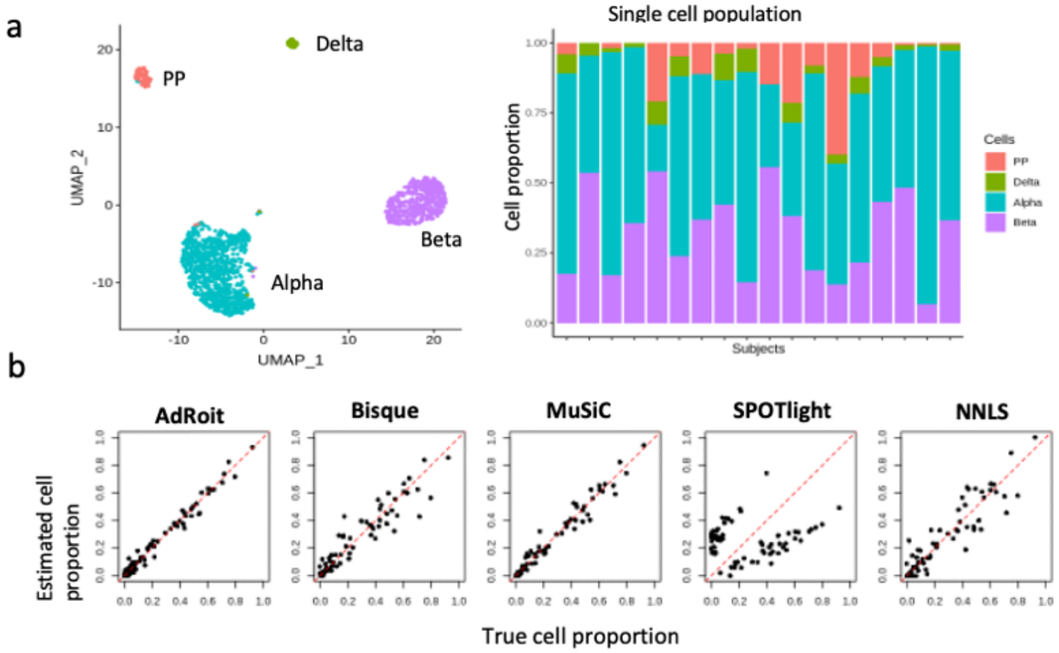


c

| | AdRoit | Bisque | MuSiC | SPOTlight | NNLS |
|---|---|---|---|---|---|
| **mAD** | 0.026 | 0.056 | 0.031 | 0.24 | 0.066 |
| **RMSD** | 0.036 | 0.082 | 0.046 | 0.25 | 0.095 |
| **Pearson** | 0.99 | 0.94 | 0.98 | 0.16 | 0.93 |
| **Spearman** | 0.97 | 0.94 | 0.98 | 0.06 | 0.91 |

**Supplementary Figure 1. Benchmark five methods on human pancreatic islets data.** a, Human islets single cell data contains 4 cell types (Alpha, Beta, PP, and Delta cells)[1] from 18 subjects.The cell proportion varies across different subjects. b, c, AdRoit achieves a leading accuracy when applied to the bulk data synthesized from the single cell data. Each dot on the scatterplot is a cell type from one subject. The reference used for deconvoluting each synthetic bulk sample excludes the cells used to synthesize that sample (i.e., leave-one-out).

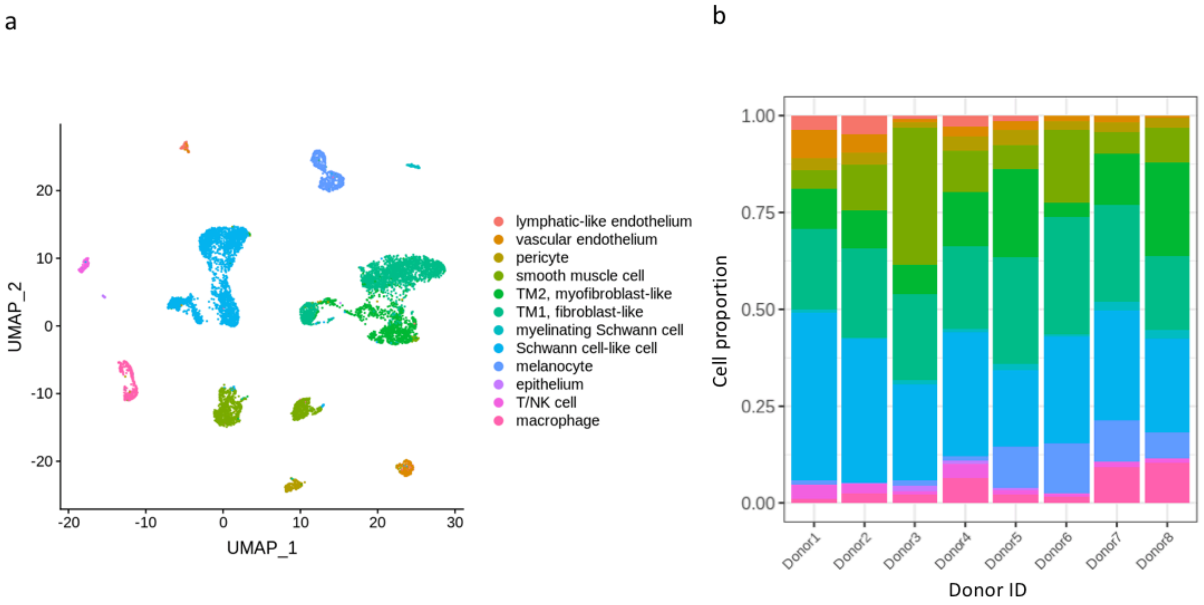


**Supplementary Figure 2. Trabecular meshwork single cell data reveals 12 cell types from 8 donors[2]. a,** UMAP[3] projection of single cells shows 12 distinct cell types. **b,** The cell proportion of each cell type varies across different donors.

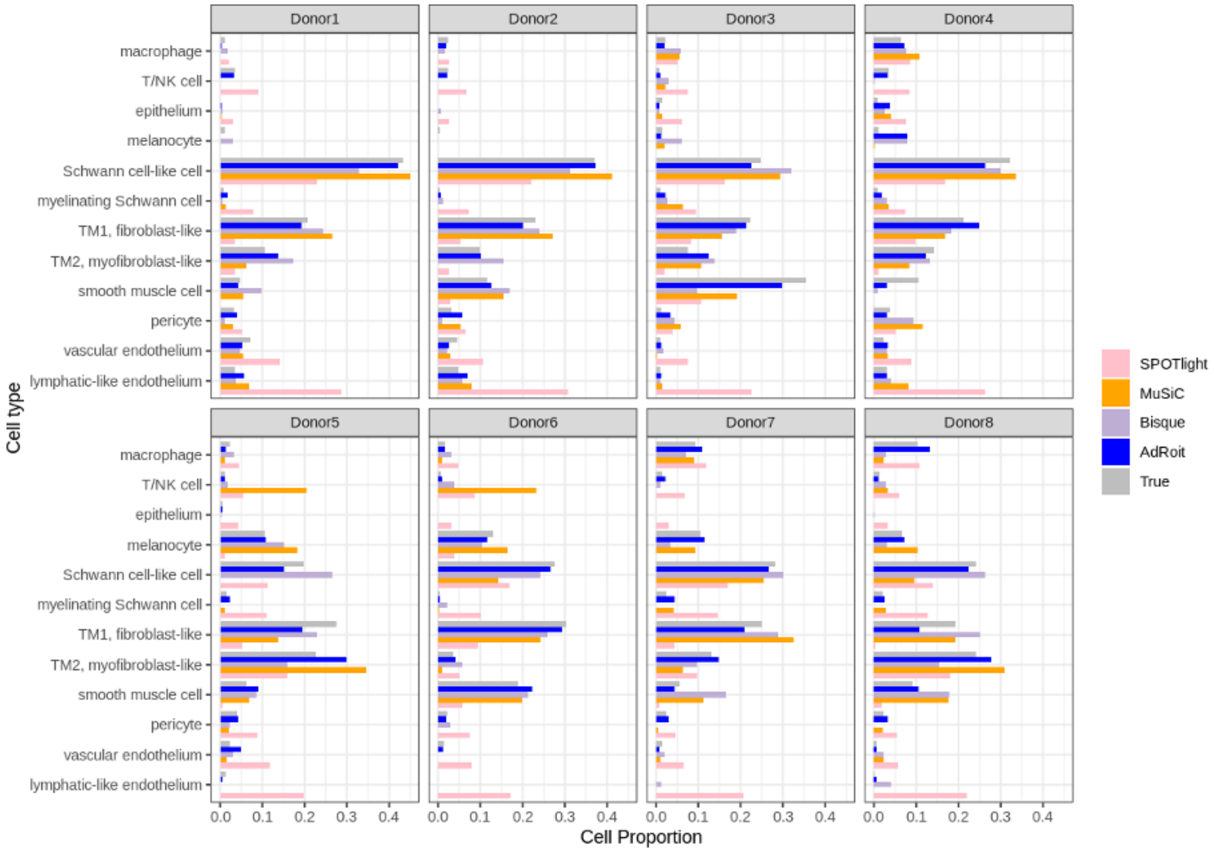


**Supplementary Figure 3. Comparison of estimated cell type proportions by AdRoit, Bisque[4], MuSiC[5] and SPOTlight[6] on the trabecular meshwork data[2] of each donor.** For each sample, AdRoit's estimates (blue bars) are more consistent with the true cell type proportions (grey bars) than the other methods.

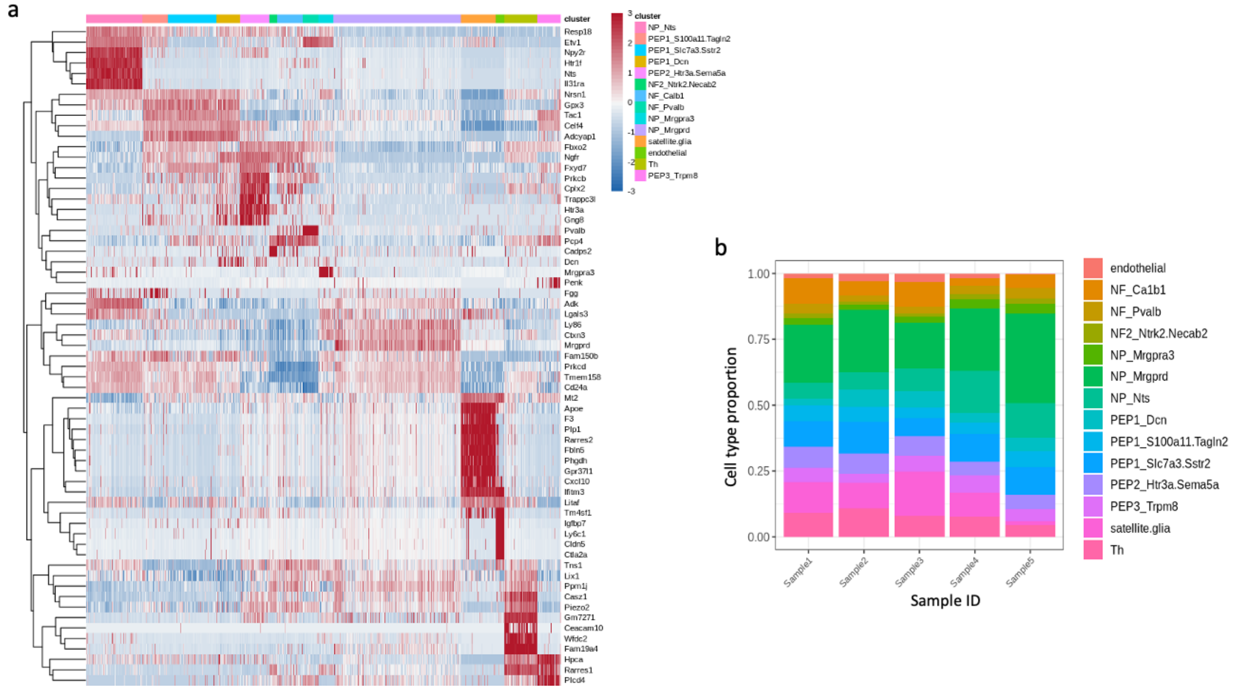


**Supplementary Figure 4. Dorsal root ganglion single cell shows 14 cell types including 3 subtypes of neurofilament-containing neurons, 3 subtypes of non-peptidergic neurons, and 5 subtypes of peptidergic neurons.** a, Heatmap of top markers shows the distinction and the similarity between the cell subtypes. b, The proportion of each cell type varies from 0.5% to 33.71% across different samples.

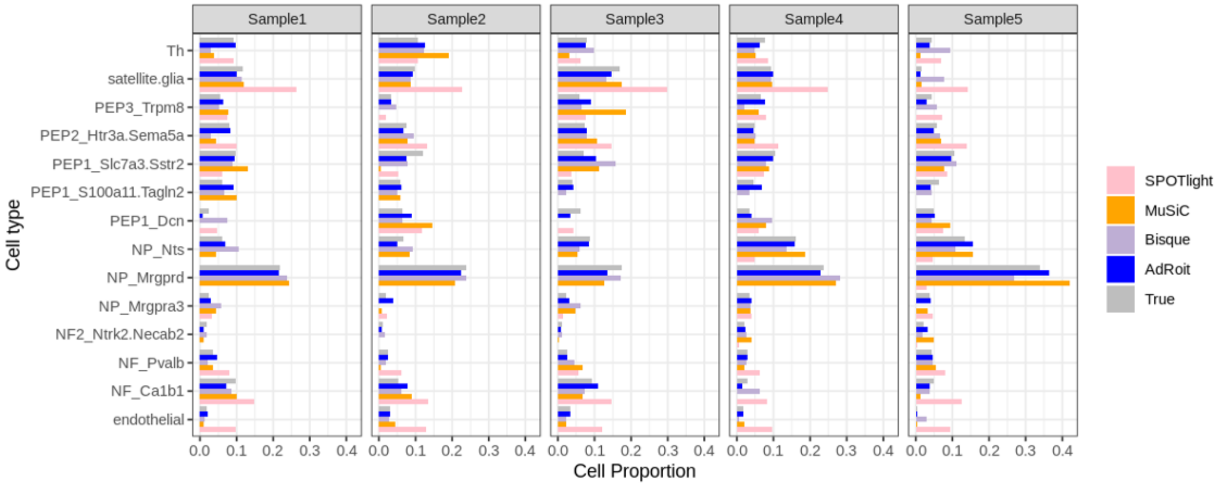


**Supplementary Figure 5. Comparison of estimated cell type proportions by AdRoit, Bisque[4], MuSiC[5] and SPOTlight[6] on the synthetic dorsal root ganglion bulk samples.** For each sample, AdRoit's estimates (blue bars) are more consistent with the true cell type proportions (grey bars) than the other methods.

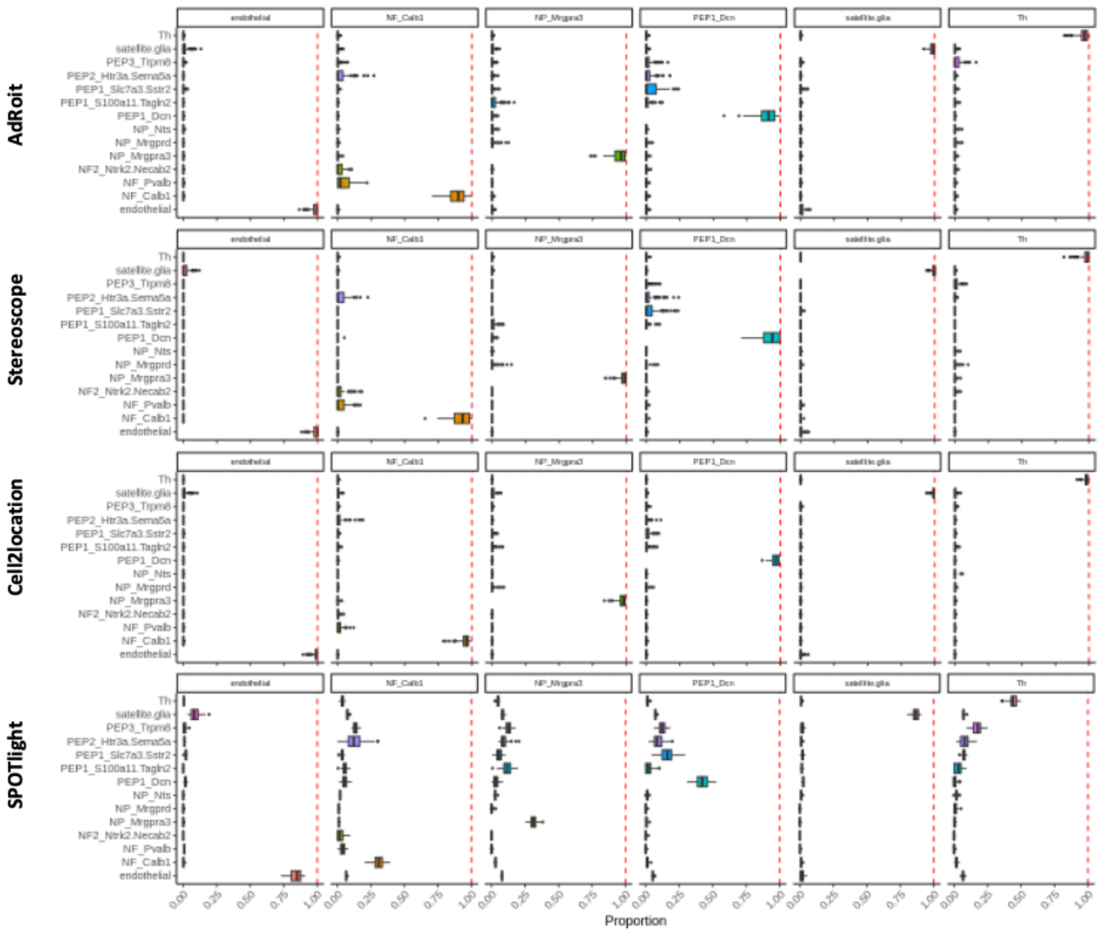


**Supplementary Figure 6. Comparing the deconvolution performance on simulated spatial spots containing a pure cell population.** All methods except SPOTlight accurately estimate the present cell type close to 1 and the absent cell types near 0. Simulations were done by sampling cells from a given cell type and adding up the UMI counts per gene. For each of the 14 DRG cell types, we repeated the simulation 100 times. The results of 6 cell types are shown here. The complete estimations for all 14 cell types are listed in Supplementary Table 11.

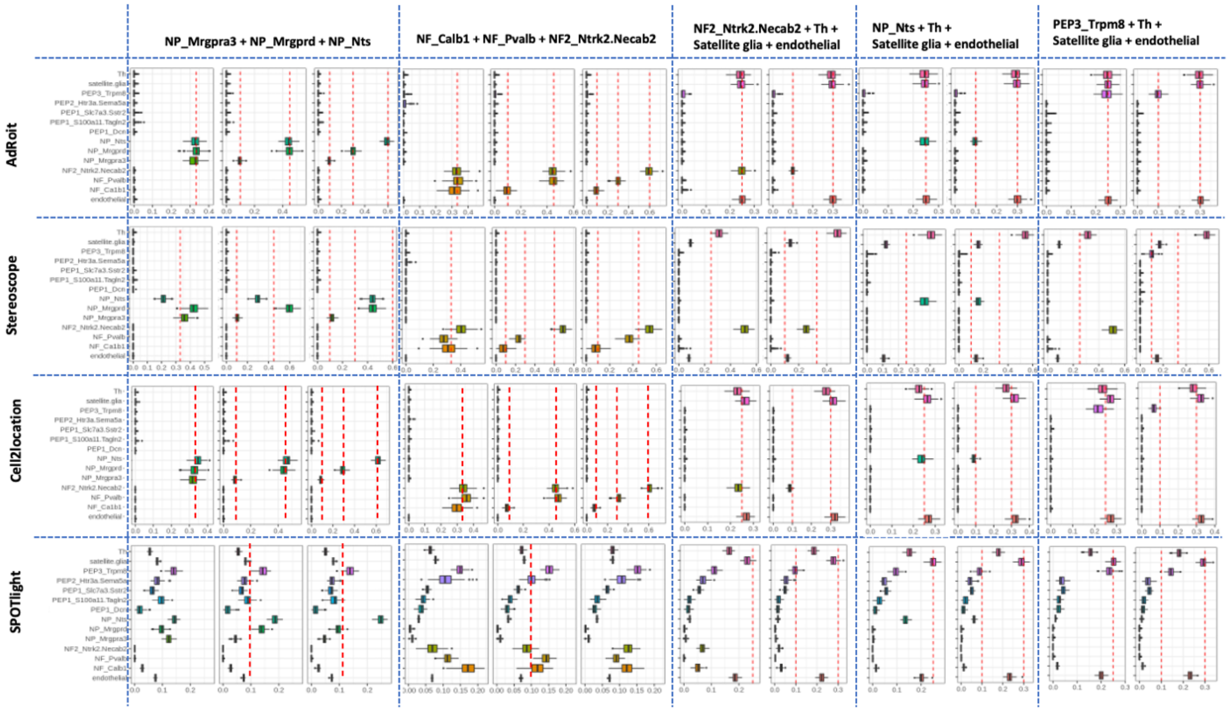


**Supplementary Figure 7. Compare deconvolution algorithms on the simulated spatial spots using additional cell mixing schemes.** 5 more sets of mixed spatial spots were simulated: 1) mixture of 3 neurofilament-containing neuron subtypes (NF); 2) mixture of 3 non-peptidergic (NP) neuron subtypes; 3) NF2_Ntrk2.Necab2 with Th, satellite glia and endothelial; 4) NP_Nts with Th, satellite glia and endothelial; and 5) PEP3_Trpm8 with Th, satellite glia and endothelial. Each simulation was repeated 100 times. Consistently for all simulation schemes, AdRoit's estimates are closer to the ground truth proportions (red lines), whereas the other methods show more deviations.

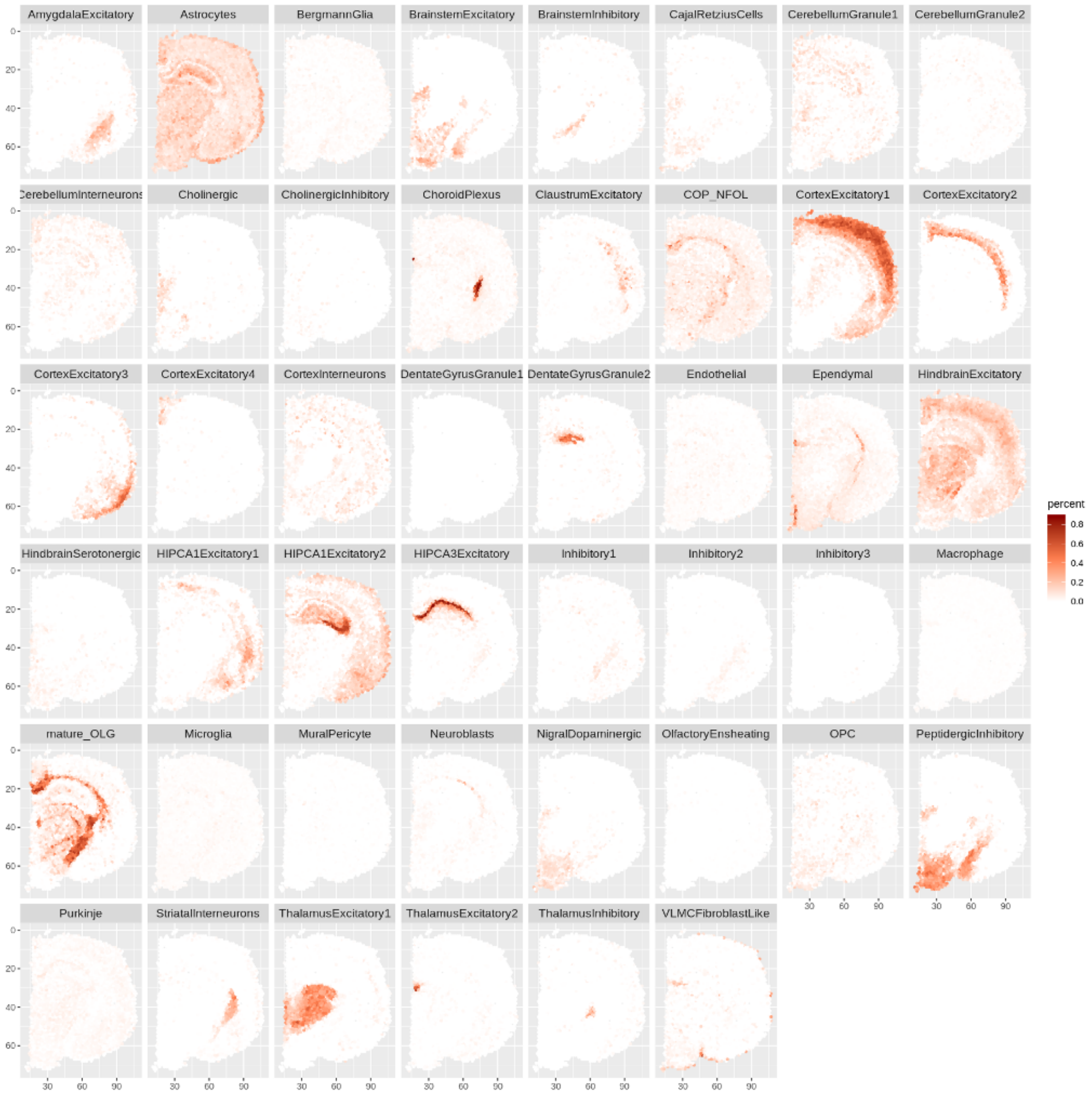


**Supplementary Figure 8. Spatial mapping of 46 brain cell types by AdRoit quantitatively depicts the content in each spot**. Spatial transcriptomics data was downloaded from 10x Genomics (see Data availability in the main text). The single cell reference data was sampled from a published mouse brain atlas[7] and consolidated into 46 cell types (see Methods in the main text).

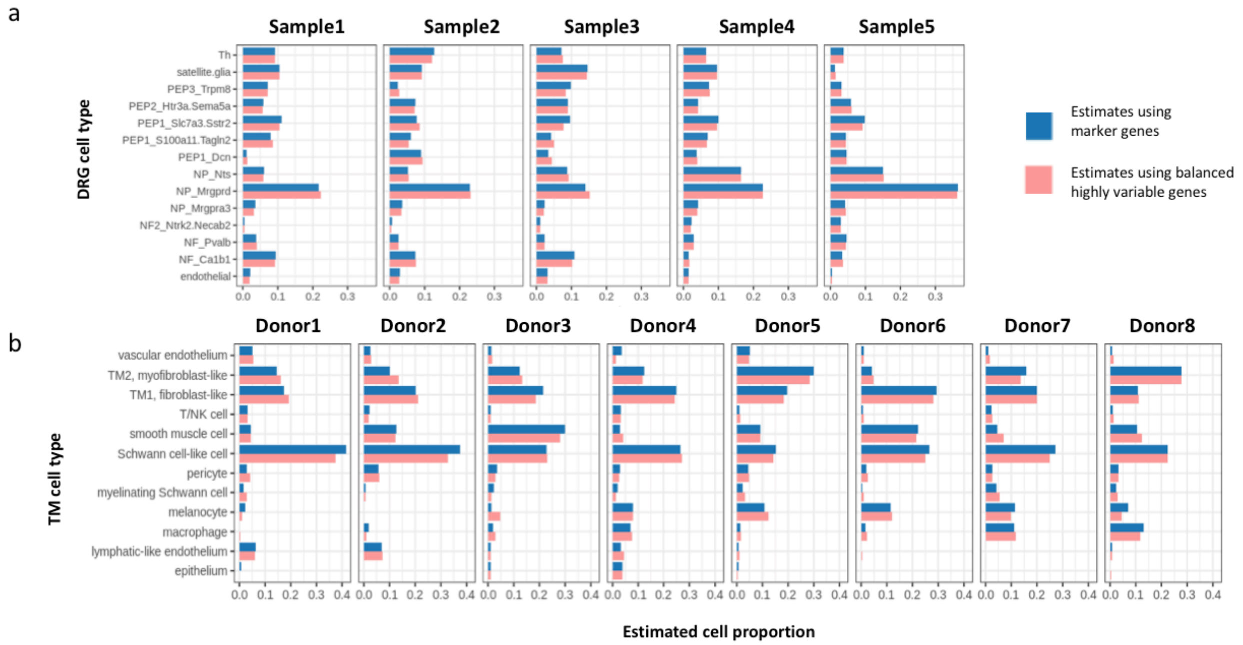


**Supplementary Figure 9. Estimation of cell proportions using either marker genes or balanced highly variable genes yields comparable results in dorsal root ganglion (DRG) data (a) and trabecular meshwork (TM) data (b).** The union of the top 200 markers (based on fold change) per cell type is compared with the top 2000 balanced highly variable genes.

### Supplementary Tables

**Supplementary Table 1. Human Islets single cell counts per subject (row).**

| | ALPHA | BETA | PP | DELTA |
|---|---|---|---|---|
| **NON_T2D_1** | 53 | 13 | 3 | 5 |
| **NON_T2D_2** | 4 | 13 | 5 | 2 |
| **NON_T2D_3** | 27 | 10 | 2 | 3 |
| **NON_T2D_4** | 14 | 10 | 3 | 0 |
| **NON_T2D_5** | 23 | 22 | 2 | 5 |
| **NON_T2D_6** | 36 | 7 | 1 | 4 |
| **NON_T2D_7** | 8 | 15 | 4 | 0 |
| **NON_T2D_8** | 14 | 16 | 9 | 3 |
| **NON_T2D_9** | 26 | 7 | 3 | 1 |
| **NON_T2D_10** | 18 | 23 | 0 | 2 |
| **NON_T2D_11** | 47 | 10 | 1 | 1 |
| **NON_T2D_12** | 84 | 48 | 0 | 2 |
| **T2D_1** | 38 | 12 | 35 | 3 |
| **T2D_2** | 50 | 18 | 10 | 5 |
| **T2D_3** | 29 | 26 | 3 | 2 |
| **T2D_4** | 138 | 135 | 2 | 5 |
| **T2D_5** | 152 | 11 | 1 | 1 |
| **T2D_6** | 125 | 76 | 1 | 5 |

**Supplementary Table 2. Leaving-one-out estimates of the cell proportions in the synthetic bulk RNA-seq data of human pancreatic islets (associated with Supplementary Fig. 1).**

| | SUBJECT.ID | PP | DELTA | ALPHA | BETA |
|---|---|---|---|---|---|
| **AdRoit** | NON_T2D_1 | 0.0254 | 0.03 | 0.7379 | 0.2067 |
| | NON_T2D_2 | 0.2524 | 0.0913 | 0.1995 | 0.4568 |
| | NON_T2D_3 | 0.056 | 0.0693 | 0.6255 | 0.2492 |
| | NON_T2D_4 | 0.1071 | 0.0067 | 0.5216 | 0.3647 |
| | NON_T2D_5 | 0.01 | 0.1801 | 0.4338 | 0.3761 |
| | NON_T2D_6 | 0.0148 | 0.0476 | 0.8236 | 0.114 |
| | NON_T2D_7 | 0.0766 | 0.0125 | 0.3083 | 0.6026 |

| | | | | | |
|---|---|---|---|---|---|
| | NON_T2D_8 | 0.2127 | 0.0892 | 0.34 | 0.358 |
| | NON_T2D_9 | 0.0992 | 0 | 0.6742 | 0.2267 |
| | NON_T2D_10 | 0.0081 | 0.0756 | 0.4713 | 0.445 |
| | NON_T2D_11 | 0.0455 | 0.0461 | 0.7173 | 0.1911 |
| | NON_T2D_12 | 0 | 0 | 0.6219 | 0.3781 |
| | T2D_1 | 0.3935 | 0.1266 | 0.3445 | 0.1354 |
| | T2D_2 | 0.1345 | 0.0959 | 0.6316 | 0.138 |
| | T2D_3 | 0.0534 | 0.057 | 0.4334 | 0.4562 |
| | T2D_4 | 0 | 0.0131 | 0.503 | 0.4838 |
| | T2D_5 | 0.0367 | 0.0086 | 0.9293 | 0.0254 |
| | T2D_6 | 0 | 0.0257 | 0.5968 | 0.3775 |
| **MuSiC** | Non_T2D_1 | 0.0255 | 0.0617 | 0.5913 | 0.3215 |
| | Non_T2D_2 | 0.158 | 0.0431 | 0.1757 | 0.6232 |
| | Non_T2D_3 | 0.0514 | 0.0723 | 0.6625 | 0.2138 |
| | Non_T2D_4 | 0.1776 | 0 | 0.4419 | 0.3805 |
| | Non_T2D_5 | 0.0188 | 0.144 | 0.4986 | 0.3386 |
| | Non_T2D_6 | 0.0008 | 0.0849 | 0.8218 | 0.0925 |
| | Non_T2D_7 | 0.1384 | 0 | 0.3002 | 0.5614 |
| | Non_T2D_8 | 0.1683 | 0.1336 | 0.3087 | 0.3894 |
| | Non_T2D_9 | 0.1598 | 0.005 | 0.6536 | 0.1817 |
| | Non_T2D_10 | 0 | 0.0712 | 0.5261 | 0.4027 |
| | Non_T2D_11 | 0.0279 | 0.0195 | 0.7426 | 0.21 |
| | Non_T2D_12 | 0 | 0.009 | 0.6107 | 0.3802 |
| | T2D_1 | 0.4083 | 0.0326 | 0.4018 | 0.1574 |
| | T2D_2 | 0.0882 | 0.0587 | 0.6537 | 0.1994 |
| | T2D_3 | 0.0561 | 0.0335 | 0.3904 | 0.52 |
| | T2D_4 | 0.0064 | 0.0165 | 0.5013 | 0.4758 |
| | T2D_5 | 0.0007 | 0.0088 | 0.9435 | 0.0471 |
| | T2D_6 | 0.0078 | 0.0128 | 0.6243 | 0.3551 |
| **NNLS** | Non_T2D_1 | 0.1468 | 0.0663 | 0.4574 | 0.3296 |
| | Non_T2D_2 | 0.2203 | 0.0379 | 0.1332 | 0.6087 |
| | Non_T2D_3 | 0 | 0.0639 | 0.6072 | 0.3289 |
| | Non_T2D_4 | 0.1198 | 0.001 | 0.4001 | 0.4791 |
| | Non_T2D_5 | 0 | 0.1826 | 0.6279 | 0.1894 |
| | Non_T2D_6 | 0 | 0.0957 | 0.8888 | 0.0155 |
| | Non_T2D_7 | 0.1291 | 0 | 0.3044 | 0.5665 |
| | Non_T2D_8 | 0.2071 | 0.1337 | 0.269 | 0.3902 |
| | Non_T2D_9 | 0.1871 | 0.004 | 0.5751 | 0.2338 |
| | Non_T2D_10 | 0.0886 | 0.0635 | 0.5124 | 0.3355 |

| | | | | | |
|---|---|---|---|---|---|
| | Non_T2D_11 | 0.2559 | 0.0191 | 0.5806 | 0.1444 |
| | Non_T2D_12 | 0 | 0.0075 | 0.6609 | 0.3316 |
| | T2D_1 | 0.5673 | 0.0345 | 0.3512 | 0.047 |
| | T2D_2 | 0.0912 | 0.0594 | 0.6201 | 0.2293 |
| | T2D_3 | 0 | 0.027 | 0.3321 | 0.6409 |
| | T2D_4 | 0 | 0.0153 | 0.6313 | 0.3534 |
| | T2D_5 | 0 | 0 | 1 | 0 |
| | T2D_6 | 0 | 0.0116 | 0.6659 | 0.3225 |
| **Bisque** | Non_T2D_1 | 0.0875 | 0.0520 | 0.4295 | 0.4309 |
| | Non_T2D_2 | 0.1246 | 0.0379 | 0.2837 | 0.5538 |
| | Non_T2D_3 | 0.0344 | 0.0553 | 0.7063 | 0.2040 |
| | Non_T2D_4 | 0.1520 | 0.0166 | 0.4737 | 0.3577 |
| | Non_T2D_5 | 0.0317 | 0.0982 | 0.5139 | 0.3561 |
| | Non_T2D_6 | 0.0000 | 0.0701 | 0.8376 | 0.0923 |
| | Non_T2D_7 | 0.1527 | 0.0175 | 0.3952 | 0.4346 |
| | Non_T2D_8 | 0.1631 | 0.1085 | 0.3878 | 0.3405 |
| | Non_T2D_9 | 0.1483 | 0.0132 | 0.6254 | 0.2131 |
| | Non_T2D_10 | 0.0678 | 0.0571 | 0.5520 | 0.3231 |
| | Non_T2D_11 | 0.1207 | 0.0268 | 0.5639 | 0.2887 |
| | Non_T2D_12 | 0.0134 | 0.0192 | 0.6001 | 0.3673 |
| | T2D_1 | 0.2695 | 0.0375 | 0.4011 | 0.2919 |
| | T2D_2 | 0.0696 | 0.0501 | 0.6079 | 0.2723 |
| | T2D_3 | 0.0352 | 0.0307 | 0.4493 | 0.4847 |
| | T2D_4 | 0.0022 | 0.0234 | 0.5974 | 0.3769 |
| | T2D_5 | 0.0000 | 0.0184 | 0.8545 | 0.1271 |
| | T2D_6 | 0.0202 | 0.0191 | 0.6669 | 0.2939 |
| **SPOTlight** | Non_T2D_1 | 0.2559 | 0.3244 | 0.3344 | 0.0853 |
| | Non_T2D_2 | 0.2669 | 0.4854 | 0.0610 | 0.1867 |
| | Non_T2D_3 | 0.2803 | 0.3372 | 0.2545 | 0.1279 |
| | Non_T2D_5 | 0.3829 | 0.2843 | 0.1745 | 0.1583 |
| | Non_T2D_6 | 0.2934 | 0.2943 | 0.3492 | 0.0631 |
| | Non_T2D_7 | 0.1964 | 0.3945 | 0.1010 | 0.3081 |
| | Non_T2D_8 | 0.2661 | 0.4689 | 0.0991 | 0.1659 |
| | Non_T2D_9 | 0.1580 | 0.4202 | 0.3206 | 0.1012 |
| | Non_T2D_10 | 0.2591 | 0.2991 | 0.2078 | 0.2340 |
| | Non_T2D_11 | 0.2080 | 0.3237 | 0.3709 | 0.0974 |
| | Non_T2D_12 | 0.1664 | 0.2776 | 0.3145 | 0.2416 |
| | Non_T2D_4 | 0.1983 | 0.4161 | 0.2062 | 0.1794 |
| | T2D_1 | 0.1676 | 0.7425 | 0.0885 | 0.0014 |

| | | | | | |
|---|---|---|---|---|---|
| | T2D_2 | 0.2591 | 0.4119 | 0.2698 | 0.0591 |
| | T2D_3 | 0.2699 | 0.3395 | 0.1633 | 0.2273 |
| | T2D_4 | 0.2095 | 0.2688 | 0.2316 | 0.2901 |
| | T2D_5 | 0.1698 | 0.3154 | 0.4937 | 0.0211 |
| | T2D_6 | 0.2065 | 0.2509 | 0.3170 | 0.2255 |

### Supplementary Table 3. Trabecular meshwork single cell counts per donor (column).

| | Donor1 | Donor2 | Donor3 | Donor4 | Donor5 | Donor6 | Donor7 | Donor8 |
|---|---|---|---|---|---|---|---|---|
| **lymphatic-like endothelium** | 33 | 44 | 10 | 19 | 17 | 0 | 0 | 1 |
| **vascular endothelium** | 66 | 41 | 10 | 15 | 27 | 26 | 18 | 5 |
| **pericyte** | 29 | 29 | 13 | 25 | 48 | 42 | 28 | 24 |
| **smooth muscle cell** | 44 | 105 | 385 | 68 | 76 | 356 | 64 | 90 |
| **TM2, myofibroblast-like** | 95 | 89 | 83 | 92 | 277 | 70 | 147 | 242 |
| **TM1, fibroblast-like** | 189 | 207 | 242 | 138 | 335 | 570 | 278 | 192 |
| **myelinating Schwann cell** | 8 | 3 | 10 | 5 | 19 | 9 | 27 | 21 |
| **Schwann cell-like cell** | 394 | 334 | 270 | 208 | 241 | 519 | 315 | 242 |
| **melanocyte** | 9 | 3 | 15 | 7 | 128 | 246 | 118 | 66 |
| **epithelium** | 2 | 0 | 15 | 6 | 6 | 1 | 0 | 0 |
| **T/NK cell** | 33 | 22 | 9 | 23 | 14 | 14 | 18 | 13 |
| **macrophage** | 9 | 22 | 24 | 42 | 28 | 31 | 102 | 103 |

### Supplementary Table 4. Leaving-one-out estimates of the cell proportions in the synthetic bulk RNA-seq data of human trabecular meshwork (associated with Fig. 2 and Supplementary Fig. 3).

| | Cell Type | Donor1 | Donor2 | Donor3 | Donor4 | Donor5 | Donor6 | Donor7 | Donor8 |
|---|---|---|---|---|---|---|---|---|---|
| **AdRoit** | lymphatic-like endothelium | 0.0601 | 0.0683 | 0.0043 | 0.0118 | 0.0025 | 0 | 0 | 0.0011 |
| | vascular endothelium | 0.0473 | 0.0177 | 0.0107 | 0.0266 | 0.0456 | 0.0073 | 0.0068 | 0.0112 |
| | pericyte | 0.0362 | 0.0538 | 0.0269 | 0.0063 | 0.0402 | 0.0136 | 0.0247 | 0.0219 |
| | smooth muscle cell | 0.0379 | 0.1182 | 0.3438 | 0.0149 | 0.0875 | 0.2282 | 0.0342 | 0.1027 |
| | TM2, myofibroblast-like | 0.122 | 0.072 | 0.0868 | 0.1171 | 0.316 | 0.0031 | 0.1228 | 0.311 |

| | | | | | | | | | |
|---|---|---|---|---|---|---|---|---|---|
| | TM1, fibroblast-like | 0.2023 | 0.232 | 0.2454 | 0.2872 | 0.1797 | 0.3254 | 0.2376 | 0.0952 |
| | myelinating Schwann cell | 0.002 | 0 | 0.0187 | 0.0447 | 0.0223 | 0.0006 | 0.0402 | 0.0223 |
| | Schwann cell-like cell | 0.4497 | 0.4038 | 0.2166 | 0.2116 | 0.1456 | 0.2559 | 0.2802 | 0.234 |
| | melanocyte | 0 | 0 | 0 | 0.0863 | 0.1286 | 0.1319 | 0.1253 | 0.0734 |
| | epithelium | 0.0043 | 0 | 0.0085 | 0.126 | 0.0053 | 0.0034 | 0 | 0 |
| | T/NK cell | 0.0342 | 0.021 | 0.0118 | 0.0301 | 0.0115 | 0.0093 | 0.0202 | 0.0095 |
| | macrophage | 0.0042 | 0.0134 | 0.0264 | 0.0374 | 0.0153 | 0.0213 | 0.108 | 0.1177 |
| **MuSiC** | lymphatic-like endothelium | 0.0701 | 0.079 | 0.0143 | 0.0805 | 0 | 0 | 0.0001 | 0 |
| | vascular endothelium | 0.0537 | 0.0291 | 0.0029 | 0.0319 | 0.0146 | 0 | 0.0112 | 0.022 |
| | pericyte | 0.0313 | 0.0537 | 0.0596 | 0.1147 | 0.0212 | 0 | 0.0063 | 0.021 |
| | smooth muscle cell | 0.0539 | 0.1551 | 0.192 | 0 | 0.0698 | 0.1975 | 0.1128 | 0.1751 |
| | TM2, myofibroblast-like | 0.0616 | 0 | 0.1074 | 0.0839 | 0.3455 | 0.0083 | 0.0631 | 0.31 |
| | TM1, fibroblast-like | 0.2658 | 0.2703 | 0.1554 | 0.1676 | 0.1368 | 0.2427 | 0.3259 | 0.1921 |
| | myelinating Schwann cell | 0.0123 | 0 | 0.0623 | 0.0352 | 0.0116 | 0.002 | 0.0412 | 0.0278 |
| | Schwann cell-like cell | 0.4486 | 0.4128 | 0.2939 | 0.3362 | 0.002 | 0.1429 | 0.2553 | 0.0957 |
| | melanocyte | 0.0001 | 0 | 0.0187 | 0.0005 | 0.1831 | 0.1642 | 0.0926 | 0.1024 |
| | epithelium | 0.0025 | 0 | 0.0162 | 0.0411 | 0 | 0 | 0 | 0 |
| | T/NK cell | 0 | 0 | 0.022 | 0 | 0.2056 | 0.2324 | 0.0008 | 0.0319 |
| | macrophage | 0 | 0 | 0.0555 | 0.1084 | 0.0099 | 0.0101 | 0.0907 | 0.022 |
| **Bisque** | lymphatic-like endothelium | 0.0379 | 0.0589 | 0.0096 | 0.0404 | 0.0020 | 0.0000 | 0.0122 | 0.0407 |
| | vascular endothelium | 0.0463 | 0.0214 | 0.0180 | 0.0306 | 0.0313 | 0.0000 | 0.0200 | 0.0221 |
| | pericyte | 0.0119 | 0.0081 | 0.0440 | 0.0923 | 0.0240 | 0.0289 | 0.0000 | 0.0000 |
| | smooth muscle cell | 0.0978 | 0.1696 | 0.0969 | 0.0097 | 0.0852 | 0.2134 | 0.1661 | 0.1778 |
| | TM2, myofibroblast-like | 0.1731 | 0.1554 | 0.1398 | 0.1331 | 0.1580 | 0.0587 | 0.0965 | 0.1546 |
| | TM1, fibroblast-like | 0.2443 | 0.2387 | 0.1891 | 0.1829 | 0.2281 | 0.2596 | 0.2899 | 0.2513 |
| | myelinating Schwann cell | 0.0051 | 0.0122 | 0.0273 | 0.0294 | 0.0000 | 0.0211 | 0.0000 | 0.0000 |
| | Schwann cell-like cell | 0.3281 | 0.3129 | 0.3195 | 0.2992 | 0.2649 | 0.2422 | 0.3016 | 0.2642 |
| | melanocyte | 0.0306 | 0.0000 | 0.0603 | 0.0790 | 0.1521 | 0.1045 | 0.0332 | 0.0305 |
| | epithelium | 0.0050 | 0.0071 | 0.0082 | 0.0267 | 0.0034 | 0.0000 | 0.0000 | 0.0011 |
| | T/NK cell | 0.0009 | 0.0000 | 0.0284 | 0.0004 | 0.0188 | 0.0395 | 0.0098 | 0.0288 |
| | macrophage | 0.0190 | 0.0156 | 0.0588 | 0.0761 | 0.0322 | 0.0319 | 0.0705 | 0.0288 |
| **SPOTlight** | lymphatic-like endothelium | 0.2873 | 0.3084 | 0.2251 | 0.2634 | 0.1987 | 0.1730 | 0.2059 | 0.2200 |
| | vascular endothelium | 0.1415 | 0.1062 | 0.0752 | 0.0882 | 0.1179 | 0.0796 | 0.0671 | 0.0564 |
| | pericyte | 0.0521 | 0.0657 | 0.0393 | 0.0531 | 0.0873 | 0.0738 | 0.0471 | 0.0544 |
| | smooth muscle cell | 0.0000 | 0.0288 | 0.1077 | 0.0000 | 0.0070 | 0.0578 | 0.0075 | 0.0171 |
| | TM2, myofibroblast-like | 0.0354 | 0.0271 | 0.0204 | 0.0110 | 0.1590 | 0.0494 | 0.0985 | 0.1799 |
| | TM1, fibroblast-like | 0.0345 | 0.0519 | 0.0836 | 0.0978 | 0.0529 | 0.0944 | 0.0439 | 0.0042 |

| | Cell type | Donor1 | Donor2 | Donor3 | Donor4 | Donor5 | Donor6 | Donor7 | Donor8 |
|---|---|---|---|---|---|---|---|---|---|
| | myelinating Schwann cell | 0.0791 | 0.0724 | 0.0958 | 0.0731 | 0.1114 | 0.1011 | 0.1448 | 0.1266 |
| | Schwann cell-like cell | 0.2291 | 0.2193 | 0.1631 | 0.1690 | 0.1136 | 0.1695 | 0.1691 | 0.1396 |
| | melanocyte | 0.0000 | 0.0000 | 0.0000 | 0.0000 | 0.0099 | 0.0377 | 0.0000 | 0.0000 |
| | epithelium | 0.0305 | 0.0254 | 0.0606 | 0.0757 | 0.0421 | 0.0306 | 0.0287 | 0.0340 |
| | T/NK cell | 0.0897 | 0.0682 | 0.0766 | 0.0826 | 0.0552 | 0.0858 | 0.0675 | 0.0599 |
| | macrophage | 0.0207 | 0.0266 | 0.0525 | 0.0861 | 0.0450 | 0.0474 | 0.1200 | 0.1079 |

**Supplementary Table 5. Estimates of cell proportions in the mixtures of myeloid and lymphoid cell types (associated with Fig. 3a).**

Provided in a separate file.

**Supplementary Table 6. Estimates of cell proportions in the bulk RNA-seq data synthesized using 6 out of the 12 trabecular meshwork cell types (associated with Fig. 3c, d). The cell types in presence are pericyte, macrophage, melanocyte, smooth muscle cell, TM1 fibroblast-like, and Schwann cell-like cells.**

| | Cell type | Donor1 | Donor2 | Donor3 | Donor4 | Donor5 | Donor6 | Donor7 | Donor8 |
|---|---|---|---|---|---|---|---|---|---|
| AdRoit | lymphatic-like endothelium | 0.0035 | 0.0052 | 0.0007 | 0.0037 | 0.0000 | 0.0000 | 0.0000 | 0.0000 |
| | vascular endothelium | 0.0046 | 0.0054 | 0.0010 | 0.0042 | 0.0000 | 0.0000 | 0.0000 | 0.0000 |
| | pericyte | 0.0469 | 0.0515 | 0.0157 | 0.0482 | 0.0594 | 0.0164 | 0.0271 | 0.0359 |
| | smooth muscle cell | 0.0653 | 0.1710 | 0.3383 | 0.0633 | 0.1185 | 0.2245 | 0.0669 | 0.1118 |
| | TM2, myofibroblast-like | 0.0150 | 0.0083 | 0.0000 | 0.0122 | 0.0383 | 0.0000 | 0.0047 | 0.0411 |
| | TM1, fibroblast-like | 0.2733 | 0.2594 | 0.2849 | 0.3282 | 0.3086 | 0.3195 | 0.2993 | 0.1994 |
| | myelinating Schwann cell | 0.0015 | 0.0000 | 0.0024 | 0.0000 | 0.0014 | 0.0000 | 0.0021 | 0.0002 |
| | Schwann cell-like cell | 0.5762 | 0.4727 | 0.2992 | 0.3845 | 0.2713 | 0.2936 | 0.3419 | 0.3342 |
| | melanocyte | 0.0000 | 0.0000 | 0.0267 | 0.0631 | 0.1835 | 0.1332 | 0.1388 | 0.0999 |
| | epithelium | 0.0004 | 0.0000 | 0.0007 | 0.0016 | 0.0000 | 0.0000 | 0.0000 | 0.0000 |
| | T/NK cell | 0.0023 | 0.0021 | 0.0012 | 0.0019 | 0.0006 | 0.0000 | 0.0000 | 0.0000 |
| | macrophage | 0.0110 | 0.0244 | 0.0290 | 0.0892 | 0.0182 | 0.0129 | 0.1192 | 0.1775 |
| MuSiC | lymphatic-like endothelium | 0.0683 | 0.0894 | 0.0155 | 0.0762 | 0.0000 | 0.0000 | 0.0000 | 0.0000 |
| | vascular endothelium | 0.0498 | 0.0245 | 0.0047 | 0.0138 | 0.0155 | 0.0000 | 0.0114 | 0.0192 |
| | pericyte | 0.0386 | 0.0484 | 0.0342 | 0.1110 | 0.0168 | 0.0000 | 0.0078 | 0.0221 |

| | | | | | | | | | |
|---|---|---|---|---|---|---|---|---|---|
| | smooth muscle cell | 0.0524 | 0.1474 | 0.2312 | 0.0003 | 0.0706 | 0.1954 | 0.1009 | 0.1521 |
| | TM2, myofibroblast-like | 0.0612 | 0.0000 | 0.1087 | 0.1083 | 0.3250 | 0.0064 | 0.0753 | 0.3036 |
| | TM1, fibroblast-like | 0.2625 | 0.2737 | 0.1608 | 0.1782 | 0.1612 | 0.2585 | 0.3096 | 0.1802 |
| | myelinating Schwann cell | 0.0105 | 0.0000 | 0.0355 | 0.0121 | 0.0104 | 0.0019 | 0.0377 | 0.0250 |
| | Schwann cell-like cell | 0.4528 | 0.4165 | 0.3160 | 0.3709 | 0.0375 | 0.1679 | 0.2697 | 0.1318 |
| | melanocyte | 0.0000 | 0.0000 | 0.0148 | 0.0004 | 0.1697 | 0.1645 | 0.0956 | 0.0901 |
| | epithelium | 0.0039 | 0.0000 | 0.0157 | 0.0311 | 0.0000 | 0.0000 | 0.0000 | 0.0000 |
| | T/NK cell | 0.0000 | 0.0000 | 0.0142 | 0.0000 | 0.1819 | 0.1937 | 0.0011 | 0.0260 |
| | macrophage | 0.0000 | 0.0000 | 0.0487 | 0.0979 | 0.0114 | 0.0118 | 0.0908 | 0.0500 |
| Bisque | lymphatic-like endothelium | 0.0249 | 0.0552 | 0.0015 | 0.0383 | 0.0000 | 0.0000 | 0.0119 | 0.0364 |
| | vascular endothelium | 0.0470 | 0.0109 | 0.0224 | 0.0476 | 0.0049 | 0.0000 | 0.0201 | 0.0235 |
| | pericyte | 0.0016 | 0.0041 | 0.0267 | 0.0711 | 0.0668 | 0.0364 | 0.0070 | 0.0145 |
| | smooth muscle cell | 0.0841 | 0.1711 | 0.1492 | 0.0078 | 0.0779 | 0.1977 | 0.1408 | 0.1834 |
| | TM2, myofibroblast-like | 0.1899 | 0.1769 | 0.1649 | 0.1395 | 0.1260 | 0.0637 | 0.0903 | 0.0862 |
| | TM1, fibroblast-like | 0.2479 | 0.2281 | 0.1720 | 0.1859 | 0.2517 | 0.2563 | 0.2892 | 0.2716 |
| | myelinating Schwann cell | 0.0040 | 0.0044 | 0.0261 | 0.0214 | 0.0133 | 0.0099 | 0.0006 | 0.0005 |
| | Schwann cell-like cell | 0.3774 | 0.3342 | 0.3043 | 0.3202 | 0.2317 | 0.2448 | 0.2975 | 0.2517 |
| | melanocyte | 0.0067 | 0.0000 | 0.0375 | 0.0652 | 0.1600 | 0.1275 | 0.0482 | 0.0406 |
| | epithelium | 0.0099 | 0.0057 | 0.0120 | 0.0206 | 0.0000 | 0.0000 | 0.0000 | 0.0000 |
| | T/NK cell | 0.0000 | 0.0000 | 0.0346 | 0.0023 | 0.0306 | 0.0399 | 0.0134 | 0.0319 |
| | macrophage | 0.0067 | 0.0095 | 0.0486 | 0.0802 | 0.0372 | 0.0238 | 0.0811 | 0.0597 |
| SPOTlight | lymphatic-like endothelium | 0.1963 | 0.2187 | 0.1981 | 0.2080 | 0.1539 | 0.1576 | 0.1913 | 0.1946 |
| | vascular endothelium | 0.1112 | 0.1120 | 0.0786 | 0.0972 | 0.0739 | 0.0727 | 0.0724 | 0.0707 |
| | pericyte | 0.0664 | 0.0710 | 0.0273 | 0.0562 | 0.1109 | 0.0668 | 0.0354 | 0.0475 |
| | smooth muscle cell | 0.0000 | 0.0307 | 0.1169 | 0.0000 | 0.0000 | 0.0475 | 0.0000 | 0.0045 |
| | TM2, myofibroblast-like | 0.0684 | 0.0622 | 0.0496 | 0.0239 | 0.1312 | 0.0840 | 0.0978 | 0.1337 |
| | TM1, fibroblast-like | 0.0410 | 0.0498 | 0.0779 | 0.1027 | 0.0673 | 0.0902 | 0.0419 | 0.0000 |
| | myelinating Schwann cell | 0.0991 | 0.0922 | 0.1042 | 0.0823 | 0.1122 | 0.1100 | 0.1336 | 0.1301 |
| | Schwann cell-like cell | 0.3299 | 0.2747 | 0.1676 | 0.2092 | 0.1484 | 0.1676 | 0.1959 | 0.1741 |
| | melanocyte | 0.0000 | 0.0000 | 0.0000 | 0.0000 | 0.0596 | 0.0498 | 0.0000 | 0.0000 |
| | epithelium | 0.0346 | 0.0287 | 0.0553 | 0.0699 | 0.0369 | 0.0342 | 0.0312 | 0.0327 |
| | T/NK cell | 0.0324 | 0.0302 | 0.0666 | 0.0397 | 0.0455 | 0.0696 | 0.0388 | 0.0397 |
| | macrophage | 0.0206 | 0.0297 | 0.0579 | 0.1111 | 0.0603 | 0.0500 | 0.1617 | 0.1723 |

**Supplementary Table 7. Sampled mouse brain single cells from Zeisel et al[7] and the consolidated cell type annotation (associated with Fig. 3c, d).**

Provided in a separate file.

**Supplementary Table 8. Estimates of cell proportions in the simulated bulk samples of high complexity (30 mouse brain cell types) (associated with Fig. 3e, f).**

Provided in a separate file.

**Supplementary Table 9. Dorsal root ganglion single cell counts per sample (column).**

| Cell type | Sample1 | Sample2 | Sample3 | Sample4 | Sample5 |
|---|---|---|---|---|---|
| **Endothelial** | 3 | 16 | 16 | 14 | 7 |
| **NF_Calb1** | 16 | 29 | 44 | 24 | 67 |
| **NF_Pvalb** | 6 | 13 | 12 | 24 | 58 |
| **NF2_Ntrk2.Necab2** | 3 | 6 | 5 | 17 | 27 |
| **NP_Mrgpra3** | 4 | 10 | 11 | 28 | 51 |
| **NP_Mrgprd** | 36 | 128 | 83 | 190 | 463 |
| **NP_Nts** | 10 | 36 | 41 | 130 | 183 |
| **PEP1_Dcn** | 4 | 35 | 29 | 28 | 67 |
| **PEP1_S100a11.Tagln2** | 10 | 31 | 19 | 36 | 85 |
| **PEP1_Slc7a3.Sstr2** | 16 | 65 | 33 | 85 | 143 |
| **PEP2_Htr3a.Sema5a** | 13 | 41 | 35 | 40 | 77 |
| **PEP3_Trpm8** | 9 | 18 | 28 | 53 | 60 |
| **Satellite.glia** | 19 | 53 | 80 | 75 | 22 |
| **Th** | 15 | 58 | 38 | 61 | 60 |

**Supplementary Table 10. Estimates of cell proportions in the synthetic bulk RNA-seq data of mouse dorsal root ganglion (associated with Fig. 4b, c and Supplementary Fig. 5).**

| | Cell type | Sample1 | Sample2 | Sample3 | Sample4 | Sample5 |
|---|---|---|---|---|---|---|
| **AdRoit** | endothelial | 0.0203 | 0.0301 | 0.0338 | 0.0164 | 0.0035 |
| | NF_Ca1b1 | 0.0723 | 0.0780 | 0.1098 | 0.0136 | 0.0369 |
| | NF_Pvalb | 0.0458 | 0.0240 | 0.0253 | 0.0288 | 0.0452 |

| | | | | | | |
|---|---|---|---|---|---|---|
| | NF2_Ntrk2.Necab2 | 0.0094 | 0.0093 | 0.0086 | 0.0232 | 0.0325 |
| | NP_Mrgpra3 | 0.0292 | 0.0388 | 0.0299 | 0.0414 | 0.0402 |
| | NP_Mrgprd | 0.2143 | 0.2234 | 0.1340 | 0.2279 | 0.3644 |
| | NP_Nts | 0.0705 | 0.0497 | 0.0858 | 0.1595 | 0.1561 |
| | PEP1_Dcn | 0.0081 | 0.0894 | 0.0351 | 0.0391 | 0.0511 |
| | PEP1_S100a11.Tagln2 | 0.0918 | 0.0626 | 0.0412 | 0.0686 | 0.0416 |
| | PEP1_Slc7a3.Sstr2 | 0.0933 | 0.0765 | 0.1053 | 0.0999 | 0.0983 |
| | PEP2_Htr3a.Sema5a | 0.0827 | 0.0675 | 0.0780 | 0.0444 | 0.0504 |
| | PEP3_Trpm8 | 0.0635 | 0.0332 | 0.0892 | 0.0763 | 0.0299 |
| | Satellite.glia | 0.1013 | 0.0915 | 0.1476 | 0.0992 | 0.0121 |
| | Th | 0.0976 | 0.1261 | 0.0765 | 0.0618 | 0.0377 |
| MuSiC | endothelial | 0.0115 | 0.0434 | 0.0240 | 0.0208 | 0.0030 |
| | NF_Ca1b1 | 0.1011 | 0.0903 | 0.0676 | 0.0000 | 0.0129 |
| | NF_Pvalb | 0.0365 | 0.0064 | 0.0679 | 0.0211 | 0.0551 |
| | NF2_Ntrk2.Necab2 | 0.0101 | 0.0000 | 0.0039 | 0.0403 | 0.0500 |
| | NP_Mrgpra3 | 0.0425 | 0.0071 | 0.0482 | 0.0366 | 0.0329 |
| | NP_Mrgprd | 0.2431 | 0.2063 | 0.1262 | 0.2701 | 0.4212 |
| | NP_Nts | 0.0433 | 0.0829 | 0.0535 | 0.1852 | 0.1560 |
| | PEP1_Dcn | 0.0000 | 0.1450 | 0.0000 | 0.0796 | 0.0949 |
| | PEP1_S100a11.Tagln2 | 0.1004 | 0.0576 | 0.0000 | 0.0000 | 0.0000 |
| | PEP1_Slc7a3.Sstr2 | 0.1318 | 0.0041 | 0.1119 | 0.0888 | 0.0769 |
| | PEP2_Htr3a.Sema5a | 0.0427 | 0.0787 | 0.1061 | 0.0484 | 0.0693 |
| | PEP3_Trpm8 | 0.0783 | 0.0000 | 0.1854 | 0.0598 | 0.0000 |
| | Satellite.glia | 0.1204 | 0.0866 | 0.1732 | 0.0973 | 0.0150 |
| | Th | 0.0383 | 0.1915 | 0.0321 | 0.0518 | 0.0128 |
| Bisque | endothelial | 0.0131 | 0.0273 | 0.0238 | 0.0067 | 0.0288 |
| | NF_Ca1b1 | 0.0874 | 0.0614 | 0.0729 | 0.0636 | 0.0381 |
| | NF_Pvalb | 0.0208 | 0.0198 | 0.0446 | 0.0249 | 0.0463 |
| | NF2_Ntrk2.Necab2 | 0.0187 | 0.0153 | 0.0108 | 0.0265 | 0.0176 |
| | NP_Mrgpra3 | 0.0567 | 0.0000 | 0.0626 | 0.0365 | 0.0015 |
| | NP_Mrgprd | 0.2392 | 0.2392 | 0.1717 | 0.2810 | 0.2681 |
| | NP_Nts | 0.1066 | 0.0919 | 0.0590 | 0.1359 | 0.1072 |
| | PEP1_Dcn | 0.0743 | 0.0637 | 0.0000 | 0.0962 | 0.0430 |
| | PEP1_S100a11.Tagln2 | 0.0674 | 0.0489 | 0.0213 | 0.0346 | 0.0427 |
| | PEP1_Slc7a3.Sstr2 | 0.0890 | 0.0779 | 0.1577 | 0.0803 | 0.1109 |
| | PEP2_Htr3a.Sema5a | 0.0307 | 0.0954 | 0.0794 | 0.0517 | 0.0652 |
| | PEP3_Trpm8 | 0.0526 | 0.0467 | 0.0656 | 0.0201 | 0.0581 |
| | Satellite.glia | 0.1147 | 0.0881 | 0.1318 | 0.0941 | 0.0776 |

| | | | | | | |
|---|---|---|---|---|---|---|
| | Th | 0.0289 | 0.1243 | 0.0990 | 0.0480 | 0.0950 |
| SPOTlight | endothelial | 0.0986 | 0.1280 | 0.1225 | 0.0958 | 0.0927 |
| | NF_Ca1b1 | 0.1490 | 0.1336 | 0.1451 | 0.0830 | 0.1237 |
| | NF_Pvalb | 0.0806 | 0.0602 | 0.0572 | 0.0636 | 0.0799 |
| | NF2_Ntrk2.Necab2 | 0.0000 | 0.0000 | 0.0000 | 0.0055 | 0.0032 |
| | NP_Mrgpra3 | 0.0334 | 0.0216 | 0.0131 | 0.0410 | 0.0474 |
| | NP_Mrgprd | 0.0000 | 0.0000 | 0.0000 | 0.0000 | 0.0285 |
| | NP_Nts | 0.0000 | 0.0000 | 0.0000 | 0.0482 | 0.0452 |
| | PEP1_Dcn | 0.0475 | 0.1164 | 0.0430 | 0.0610 | 0.0744 |
| | PEP1_S100a11.Tagln2 | 0.0000 | 0.0000 | 0.0000 | 0.0000 | 0.0000 |
| | PEP1_Slc7a3.Sstr2 | 0.0610 | 0.0534 | 0.0355 | 0.0732 | 0.0846 |
| | PEP2_Htr3a.Sema5a | 0.1002 | 0.1303 | 0.1461 | 0.1141 | 0.1390 |
| | PEP3_Trpm8 | 0.0739 | 0.0205 | 0.0765 | 0.0807 | 0.0707 |
| | Satellite.glia | 0.2644 | 0.2284 | 0.2990 | 0.2479 | 0.1416 |
| | Th | 0.0912 | 0.1075 | 0.0621 | 0.0860 | 0.0693 |

**Supplementary Table 11. Estimated cell proportions of 3,200 spatial transcriptome spots simulated by sampling and pooling multiple cell types from the mouse dorsal root ganglion single cell data (associated with Fig. 5a, Supplementary Fig. 6 and 7).**

Provided in a separate file.

**Supplementary Table 12. Estimated cell percentages of 6,000 spatial transcriptome spots simulated to evaluate the accuracy and sensitivity in deconvoluting rare cell types (associated with Fig. 5b, c).**

Provided in a separate file.

**Supplementary Table 13. Read count per gene from the 70 real bulk RNA-seq samples of human pancreatic islets and the donor information[1,8,9].**

Provided in a separate file.

**Supplementary Table 14. AdRoit-estimated cell proportions in the 70 real bulk RNA-seq samples of human pancreatic islets (associated with Fig. 6a, b and c).**

| Sample.ID | Donor.ID | ALPHA | BETA | PP | DELTA |
|---|---|---|---|---|---|
| **694267** | HS_001 | 0.2533 | 0.6473 | 0.0421 | 0.0574 |
| **728211** | HS_006 | 0.4205 | 0.4659 | 0.0548 | 0.0588 |
| **740494** | HS_008 | 0.1683 | 0.7372 | 0.0512 | 0.0434 |
| **740493** | HS_009 | 0.2239 | 0.7191 | 0.0323 | 0.0247 |
| **761866** | HS_011 | 0.1975 | 0.6676 | 0.0614 | 0.0735 |
| **761864** | HS_012 | 0.2481 | 0.6257 | 0.0725 | 0.0537 |
| **761865** | HS_013 | 0.1778 | 0.6495 | 0.0626 | 0.1101 |
| **776243** | HS_014 | 0.2694 | 0.6202 | 0.0417 | 0.0687 |
| **776244** | HS_016 | 0.2191 | 0.5926 | 0.0763 | 0.1120 |
| **782323** | HS_017 | 0.3360 | 0.5432 | 0.0597 | 0.0611 |
| **782324** | HS_017 | 0.3438 | 0.5621 | 0.0526 | 0.0415 |
| **782325** | HS_017 | 0.3612 | 0.5554 | 0.0500 | 0.0333 |
| **782326** | HS_017 | 0.3767 | 0.5434 | 0.0479 | 0.0319 |
| **782327** | HS_017 | 0.2545 | 0.5823 | 0.0662 | 0.0969 |
| **782330** | HS_018 | 0.3352 | 0.6113 | 0.0319 | 0.0215 |
| **782328** | HS_018 | 0.3283 | 0.6049 | 0.0355 | 0.0313 |
| **782329** | HS_018 | 0.3288 | 0.6164 | 0.0338 | 0.0210 |
| **797002** | HS_019 | 0.2451 | 0.7111 | 0.0269 | 0.0169 |
| **815585** | HS_022 | 0.5383 | 0.3878 | 0.0433 | 0.0307 |
| **815584** | HS_024 | 0.2223 | 0.6984 | 0.0392 | 0.0401 |
| **826369** | HS_028 | 0.2652 | 0.5625 | 0.0626 | 0.1098 |
| **826368** | HS_028 | 0.2841 | 0.5525 | 0.0602 | 0.1031 |
| **826370** | HS_029 | 0.2368 | 0.6651 | 0.0579 | 0.0401 |
| **826371** | HS_029 | 0.2555 | 0.6622 | 0.0510 | 0.0313 |
| **842228** | HS_031 | 0.1609 | 0.7688 | 0.0349 | 0.0355 |
| **842229** | HS_031 | 0.1550 | 0.7673 | 0.0370 | 0.0407 |
| **863437** | HS_037 | 0.3947 | 0.4973 | 0.0559 | 0.0521 |
| **863438** | HS_037 | 0.4446 | 0.4699 | 0.0475 | 0.0380 |
| **870429** | HS_038 | 0.2406 | 0.6039 | 0.0490 | 0.1065 |
| **870428** | HS_038 | 0.2392 | 0.6090 | 0.0487 | 0.1031 |
| **878688** | HS_039 | 0.1915 | 0.7710 | 0.0375 | 0 |
| **878689** | HS_039 | 0.2063 | 0.7541 | 0.0396 | 0 |

| | | | | | |
|---|---|---|---|---|---|
| **878691** | HS_040 | 0.1956 | 0.7425 | 0.0490 | 0.0128 |
| **878690** | HS_040 | 0.1904 | 0.7421 | 0.0501 | 0.0174 |
| **881713** | HS_041 | 0.1000 | 0.8352 | 0.0323 | 0.0325 |
| **881714** | HS_041 | 0.0995 | 0.8316 | 0.0352 | 0.0337 |
| **883377** | HS_042 | 0.2144 | 0.7061 | 0.0385 | 0.0410 |
| **883378** | HS_042 | 0.2071 | 0.7069 | 0.0394 | 0.0466 |
| **884844** | HS_043 | 0.2778 | 0.6160 | 0.0469 | 0.0593 |
| **884845** | HS_043 | 0.3033 | 0.5944 | 0.0471 | 0.0552 |
| **887299** | HS_044 | 0.2320 | 0.6928 | 0.0360 | 0.0392 |
| **887298** | HS_044 | 0.2321 | 0.6947 | 0.0356 | 0.0376 |
| **892241** | HS_045 | 0.2864 | 0.6809 | 0.0283 | 0.0044 |
| **892242** | HS_045 | 0.2818 | 0.6808 | 0.0285 | 0.0088 |
| **892243** | HS_046 | 0.2818 | 0.6176 | 0.0317 | 0.0689 |
| **892244** | HS_046 | 0.2769 | 0.6261 | 0.0317 | 0.0653 |
| **894380** | HS_047 | 0.1254 | 0.7969 | 0.0432 | 0.0345 |
| **894379** | HS_047 | 0.1490 | 0.7737 | 0.0445 | 0.0328 |
| **896597** | HS_048 | 0.2821 | 0.6839 | 0.0307 | 0.0033 |
| **896598** | HS_048 | 0.2849 | 0.6796 | 0.0327 | 0.0028 |
| **898041** | HS_049 | 0.1282 | 0.7833 | 0.0528 | 0.0357 |
| **898042** | HS_049 | 0.1357 | 0.7699 | 0.0590 | 0.0354 |
| **903919** | HS_050 | 0.2805 | 0.6390 | 0.0409 | 0.0396 |
| **903920** | HS_050 | 0.2418 | 0.6747 | 0.0417 | 0.0418 |
| **903922** | HS_051 | 0.3128 | 0.6487 | 0.0384 | 0 |
| **903921** | HS_051 | 0.3093 | 0.6440 | 0.0417 | 0.0050 |
| **906490** | HS_052 | 0.4333 | 0.4354 | 0.0575 | 0.0738 |
| **906491** | HS_052 | 0.4425 | 0.4275 | 0.0581 | 0.0719 |
| **906493** | HS_053 | 0.2047 | 0.7478 | 0.0334 | 0.0141 |
| **906492** | HS_053 | 0.1909 | 0.7585 | 0.0331 | 0.0175 |
| **942670** | HS_063 | 0.4303 | 0.5165 | 0.0316 | 0.0216 |
| **942671** | HS_063 | 0.3936 | 0.5495 | 0.0319 | 0.0249 |
| **958778** | HS_066 | 0.2321 | 0.6512 | 0.0514 | 0.0654 |
| **958779** | HS_066 | 0.2340 | 0.6618 | 0.0498 | 0.0544 |
| **974251** | HS_068 | 0.3218 | 0.6184 | 0.0531 | 0.0068 |
| **974252** | HS_068 | 0.3208 | 0.6208 | 0.0511 | 0.0073 |
| **976624** | HS_069 | 0.2714 | 0.5616 | 0.0778 | 0.0893 |
| **976623** | HS_069 | 0.2701 | 0.5622 | 0.0782 | 0.0895 |
| **1001636** | HS_076 | 0.3030 | 0.5590 | 0.0523 | 0.0857 |
| **1001637** | HS_076 | 0.2782 | 0.5682 | 0.0553 | 0.0983 |

**Supplementary Table 15. The cell type composition in the pancreatic islets of 4 subjects (rows) revealed by RNA fluorescence in situ hybridization[1] (associated with Fig. 6b).**

| | GCG(ALPHA) | INS(BETA) | PPY(PP) | SST(DELTA) |
|---|---|---|---|---|
| **HS_022** | 58.90% | 33.20% | 3.00% | 4.80% |
| **HS_024** | 33.40% | 58.90% | 1.40% | 6.30% |
| **HS_028** | 25.70% | 59.60% | 9.80% | 4.90% |
| **HS_029** | 34.60% | 53.60% | 7.20% | 4.60% |

**Supplementary Table 16. AdRoit-estimated cell proportions in the real mouse brain spatial transcriptome spots (associated with Fig. 6d and Supplementary Fig. 8).**

Provided in a separate file.
